# High-Throughput Epigenetic Profiling Immunoassays for Accelerated Disease Research and Clinical Development

**DOI:** 10.1101/2024.12.05.626944

**Authors:** Priscilla Van den Ackerveken, Clotilde Hannart, Dorian Pamart, Robin Varsebroucq, Marion Wargnies, Olivia Thiry, Marie Lurkin, Séverine Vincent, Muriel Chapelier, Guillaume Rommelaere, Marielle Herzog

## Abstract

Epigenetics, which examines the regulation of genes without modification of the DNA sequence, plays a crucial role in various biological processes and disease mechanisms. Among the different forms of epigenetic modifications, histone post-translational modifications (PTMs) are important for modulating chromatin structure and gene expression. Aberrant levels of histone PTMs are implicated in a wide range of diseases, including cancer, making them promising targets for biomarker discovery and therapeutic intervention. In this context, blood, tissues, or cells serve as valuable resources for epigenetic research and analysis.

Traditional methods such as mass spectrometry and western blotting are widely used to study histone PTMs, providing qualitative and (semi)quantitative information. However, these techniques often face limitations that could include throughput and scalability, particularly when applied to clinical samples. To overcome these challenges, we developed and validated 13 Nu.Q^®^ immunoassays to detect and quantify specific histone PTM-nucleosomes from K2EDTA plasma samples. Then, we tested these assays on other types of samples, including chromatin extracts from frozen tissues, as well as cell lines and white blood cells

Our findings demonstrate that the Nu.Q^®^ assays offer high specificity, sensitivity, precision and linearity, making them effective tools for epigenetic profiling. A comparative analysis of HeLa cells using mass spectrometry, Western blot, and Nu.Q^®^ immunoassays revealed a consistent histone PTMs signature, further validating the effectiveness of these assays. Additionally, we successfully applied Nu.Q^®^ assays across various biological samples, including human tissues from different organs and specific white blood cell subtypes, highlighting their versatility and applicability in diverse biological contexts.

## Introduction

Within the nucleus of eukaryotic cells, genomic DNA is intricately organized into a highly condensed structure known as chromatin. The fundamental unit of chromatin is the nucleosome, which consists of an octamer of core histone proteins (H2A, H2B, H3, and H4) around which approximately 147 base pairs of DNA are tightly wound (1–3). These nucleosomes are connected by stretches of linker DNA and are often stabilized by linker histones such as H1, which further contribute to chromatin’s hierarchical structure (4, 5). Given that all cells within an organism have identical DNA sequence, epigenetic regulation is essential for providing each cell its unique identity and specialized function, known as cell-type-specific epigenetic signatures (6, 7). Among the various mechanisms of epigenetic regulation, nucleosome positioning, histone variants and histone post-translational modifications (PTMs) play a central role by modulating chromatin structure (8–11) and are vital for defining and preserving cell identity (7, 12). Disruptions in these mechanisms can lead to disturbances in chromatin organization and gene expression, contributing to pathological conditions and disease development (13–16). Notably, cancer is characterized by these epigenetic changes, which are considered as a hallmark of this disease (17–22). As a result, the altered epigenome of tumor cells is increasingly recognized as potentially effective biomarkers for detecting cancer conditions and classifying tumor types (23–26). Moreover, since such disruptions may be reversible and can be modulated with epigenetic drug treatments (27–31), monitoring epigenetic marks has emerged as a promising target for evaluating cancer therapies, as well as to establish personalized medicine (32–35). Despite significant advances in understanding the key molecular pathways involved in cancer, further research is still necessary to enhance our understanding of the epigenetic signature of tumor cells and to bridge the gap from preclinical models to clinical applications. In this context, patient-derived tissues obtained from surgery or tumor biopsies provide a valuable and renewable source of material for preclinical analyses, offering an opportunity to directly study tumor-specific epigenetic changes and their clinical relevance. Additionally, isolated peripheral blood mononuclear cells (PBMCs) can inform insights into the underlying mechanisms of the immune response, establishing a complementary model for preclinical research in both cancer (36–38) or other diseases (39, 40).

Several advanced tools are available to characterize epigenetic marks. Chromatin immunoprecipitation (ChIP), combined with either sequencing (ChIP-Seq) or PCR (ChIP-PCR), are invaluable for mapping histone modifications at specific genomic loci, using antibodies to pull down DNA sequences associated with particular histone PTMs and unraveling the intricate relationships between histone modifications and gene regulation(41, 42). Mass spectrometry (MS) provides a complementary approach by enabling the comprehensive analysis of PTMs across the proteome, offering detailed insights into the presence and abundance of these modifications in various cellular contexts (43, 44). However, these methods have limitations, particularly when applied to clinical samples. The low abundance of certain PTMs, along with the complexity of peptide mixtures, often necessitates further validation using additional techniques like Western blot (WB) and immunohistochemistry (IHC). While WB and IHC are useful for detecting and visualizing specific histone PTMs, their low throughput limits their utility in large-scale studies and clinical practice. Given these challenges, immunoassays have emerged as a promising alternative for the study of histone PTMs. These assays offer the opportunity of a reliable and quantitative measure of circulating nucleosomes and their associated epigenetic modifications, combining ease of use with the capacity for high-throughput analysis. Immunoassays could thus facilitate comprehensive profiling of histone PTM patterns and are particularly well-suited for clinical applications, especially for tracking epigenetic patterns in blood (45–48).

In this study, we have analytically validated 13 Nu.Q^®^ tests in blood for the detection of specific histone PTM-nucleosomes (i.e. 12 Histone-PTMs and 1 Histone-mutation) in K2EDTA plasma samples, demonstrating their performance in terms of specificity, sensitivity, precision and linearity. We then conducted a comparative analysis of chromatin extracts from HeLa cells using MS, WB, and the Nu.Q^®^ immunoassays. Our findings revealed a consistent histone PTMs signature across all three techniques, underscoring the robustness of these Nu.Q^®^ assays. . Additionally, we extended our analysis to chromatin extracts from frozen human tissues of various organ origins, successfully showcasing the versatility of the Nu.Q^®^ assays in different biological contexts. These assays were further applied to specific white blood cell subtypes, highlighting their effectiveness in diverse epigenetic landscapes. Overall, our study demonstrates that the analytically validated Nu.Q^®^ assays represent a significant advancement in high-throughput epigenetic profiling. These tools have the potential to accelerate disease research and clinical development, offering a transformative asset in the field of epigenetics. Altogether, by analyzing tissue and white blood cells, researchers can leverage these epigenetic tools to gain profound insights into cancer development and identify key therapeutic targets, with the ultimate goal of advancing the field of personalized medicine.

## Results

### Nu.Q^®^ Histone PTMs assays are analytically validated chemiluminescent immunoassays that allow a specific, precise quantification of histone-PTMs in K2EDTA plasma samples

To enable robust and reproducible quantification of histone-PTMs associated to circulating nucleosomes (PTMs-nucleosome) in K2EDTA plasma samples, 13 automated chemiluminescent immunoassays (ChLIA) have been developed. Those sandwich immunoassays (Nu.Q^®^) used an anti-histone PTMs capture antibody coated on a solid phase, consisting of magnetic particles, combined with a conformational anti-nucleosome antibody coupled with acridinium ester molecules in detection (Supporting Figure1). The analytical performance of the 13 immunoassays was rigorously evaluated in K2EDTA plasma samples. First, we assessed the specificity of each assay using recombinant nucleosomes (rNucl.) that exhibit specific histone PTMs, histone variants, or the H3K27M mutation. The Nu.Q^®^ assays demonstrate minimal/no cross-reactivity, (ranging from 0% to 3%) with most assays (Figure 1A). An exception is the Nu.Q^®^ H3K27Ac assay, which exhibits a slight cross-reactivity with the rNucl. H3K27M (8% cross-reactivity). Additionally, the results confirm that the Nu.Q^®^ H3.1 assay recognizes all recombinant PTMs-H3.1 nucleosomes, except the H3.3 variant, with 0% cross-reactivity observed. The precision of the 13 different Nu.Q^®^PTMs assays was then assessed by determining the within-run (Intra-CV) and within-laboratory (Inter-CV) coefficients of variation. The Intra-CV values range from 0.9% to 7.6%, while the Inter-CV values vary from 1.4% to 16.1% (Figure 1B), indicating high consistency and reliability of the assays. Then, the sensitivity of the 13 assays was evaluated, demonstrating good detection performance with a LOB ranging from 0.10 to 1.90 ng/mL; a LOD from 0.80 to 3.57ng/mL and a LOQ from 0.8 to 8.1 ng/mL defined as the assay limit (Figure 1C). Additionally, we determined the linear range of each Nu.Q^®^PTMs assay using K2EDTA plasma samples (Figure 1D), and the linearity of the assay curves was confirmed, with R² values ranging from 0.9985–1.00 across all assays. Overall, these analytical performances confirm that the 13 Nu.Q^®^PTMs ChLIA assays are highly precise and specific for the measurement of histone PTMs and histone mutation-nucleosome concentrations in plasma samples.

**Figure 1.**
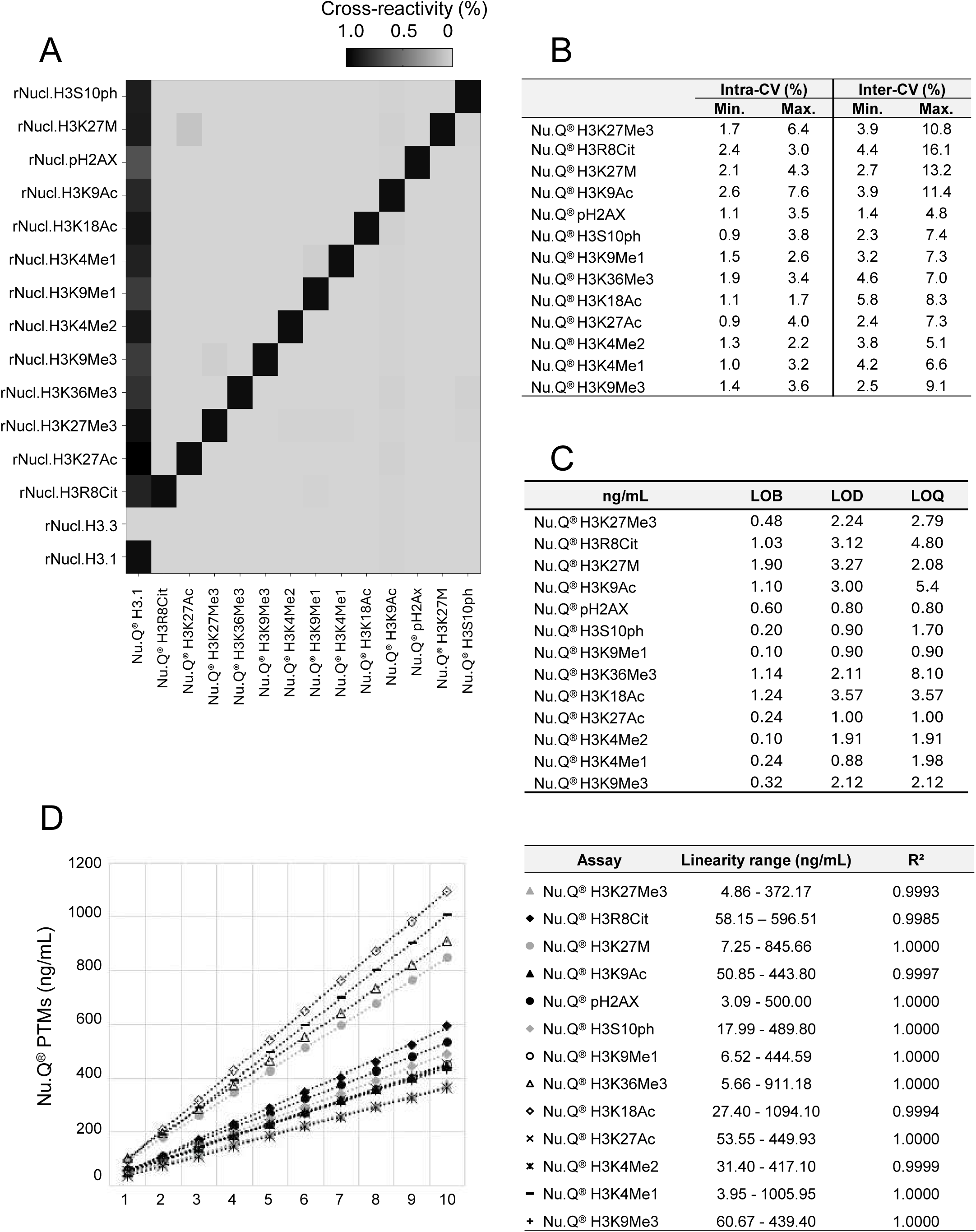
Analytical validation of Nu.Q^®^ Histone PTMs Chemiluminescent Immunoassays (ChLIA). **A.** The heat map illustrates the specificity of Nu.Q^®^H3.1 and 13 Nu.Q^®^PTMs chemiluminescent immunoassays tested against recombinant nucleosomes (rNucl.) with various histone post-translational modifications, histone variants, and the H3K27M mutation. Results show the % of cross-reactivity among the Nu.Q^®^ assays. **B.** Min. and Max. of the Intra-assay (Intra-CV) and inter-assay (Inter-CV) coefficients of variation for each of the 13 Nu.Q^®^PTMs assays. **C.** Limits of blank (LOB), limits of detection (LOD) and limits of quantification (LOQ) estimated for all Nu.Q^®^PTMs assays. **D.** Linearity range of the assays and the related R² values for all curves.

### Nu.Q^®^ Histone PTMs assays, Mass Spectrometry and Western Blotting show consistent outcomes in cell line studies

The profile of histone PTMs present on the N-terminal tails of Histone H3 in HeLa cell chromatin extracts was first characterized using MS. Through this approach, 29 different histone proteoforms were identified allowing the characterization of 17 distinct histone PTMs located at nine different sites as well as two histone H3 variants (H3.1/H3.2 and H3.3). MS analysis revealed that unmodified and methylated histone H3 peptides were more abundant, whereas acetylated histone H3 peptides were present at much lower levels in these extracts (Figure 2A). To further explore the histone H3 PTMs pattern in these cell-derived chromatin extracts, we performed Western blot analysis, which corroborated the MS findings. The WB results demonstrated strong signal intensities for methylated marks on H3 (H3K4Me1, H3K9Me1, H3K9Me3, H3K27Me3, H3K36Me3) and weaker signals for acetylation marks (H3K9Ac, H3K18Ac, H3K27Ac) and citrullination marks (H3R8Cit) (Figure 2B and Supporting Figure 2). We then quantified the histone PTMs-nucleosome pattern using Nu.Q^®^ immunoassays, which further confirmed the epigenetic profile observed by MS and WB (Figure 2C). Indeed, the Nu.Q^®^ assays results revealed low levels of acetylated histone H3.1 nucleosomes (n=4; mean±SD : H3K9Ac= 0.81±0.10%, H3K18Ac= 0.52±0.10%, H3K27Ac= 0.21±0.04%) and H3R8Cit-nucleosomes (n=4; mean±SD: H3R8Cit= 1.84±0.43%). In contrast, higher proportions of methylated histone H3.1 nucleosomes were observed (n=4; mean±SD: H3K4Me1= 10.46±1.42%; H3K4Me2= 3.34±0.56%; H3K9Me1= 21.49±3.62%; H3K9Me3= 40.88±4.07%; H3K27Me3= 31.04±2.16% and H3K36Me3= 25.39±1.87%). Given the low endogenous expression of acetylated histone H3 in HeLa cells, we aimed to assess whether the immunoassays are effective in detecting acetylation markers under induced acetylation conditions. For that purpose, HeLa cells were treated with sodium butyrate (NaB), a histone deacetylase (HDAC) inhibitor, and the levels of three acetylated nucleosomes and two methylated nucleosomes were quantified using Nu.Q^®^ immunoassays. As expected, NaB treatment led to a significant upregulation of all tested acetylated H3.1 nucleosomes ratio (n=3; mean±SD untreated vs NaB-treated : H3K9Ac= 0.88±0.69% vs 85.08±18.20% (**: *p*□<□*0.01)*; H3K18Ac= 1.35±1.42% vs 109.22±10.85% (****: *p*□<□*0.0001)*; H3K27Ac= 0.33±0.54% vs 22.11±8.02% (**: *p*□<□*0.01)*) while methylated H3.1 nucleosome ratio levels remained relatively constant (n=3: mean±SD untreated vs NaB-treated: H3K27Me3= 18.86±11.94% vs 31.08±5.86% (ns: *p*=0.1867); H3K9Me3= 25.88±17.37% vs 35.38±7.06% (ns: *p* =□*0.4299)*) (Figure 3A). Then, to further demonstrate the capacity of the immunoassays to assess histone PTMs changes in response to an epigenetic treatment, HeLa cells were exposed to GSK343 drugs, an EZH2 inhibitor, and the level of PTMs-nucleosomes were quantified using Nu.Q^®^ immunoassays (Figure 3B). As anticipated, GSK343 treatment led to a significant downregulation of H3K27Me3-nucleosomes ratio (n=3; mean±SD untreated vs GSK343-treated : 22.43±2.05% vs 13.95±2.06% (**: *p*□<□*0.01*) while other methylated or Acetylated H3.1 nucleosome ratio levels remained constant (n=3: mean±SD untreated vs GSK343-treated: H3K9Me3= 27.84±3.71% vs 24.75±0.20% (ns: *p*=0.2236); H3K18Ac= 0.48±0.05% vs 0.56±0.19% (ns: *p* =0.508); H3K9Ac= 0.83±0.16% vs 0.83±0.10% (ns: *p* =0.9994); H3K27Ac= 0.28±0.09% vs 0.34±0.06% (ns: *p* =0.3211)). Altogether, the results demonstrate the capability of Nu.Q^®^ immunoassays to accurately characterize histone PTMs in cell lines, delivering results consistent with those obtained through MS and WB.

**Figure 2.**
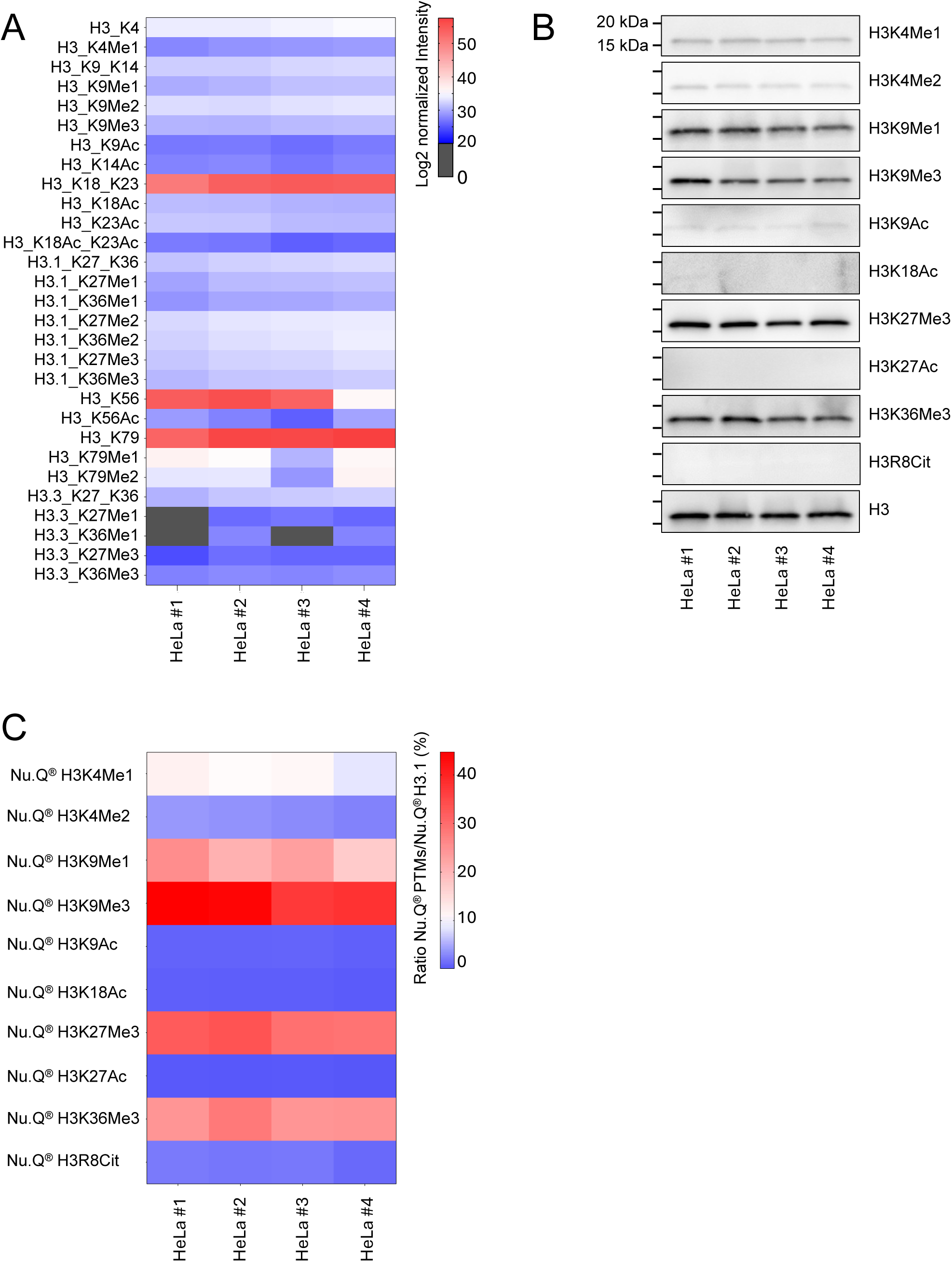
Histone-PTMs profile present in the N-terminal tails of histone H3 from HeLa cells chromatin extracts as defined by Mass Spectrometry (LC-MS/MS), Western Blotting (WB), and Nu.Q^®^ Histone-PTMs assays. **A.** The heat map illustrates the relative normalized intensity of histone PTMs detected by LC-MS/MS. Normalized raw intensities are represented on a log_2_ scale, with each cell corresponding to a specific histone PTM across four different HeLa cell chromatin extracts (HeLa #1-4). Gray cells indicate undetected histone-PTMs. **B.** Western blot analysis targeting 10 different PTMs of the histone H3 N-terminal tail with a representative result for Histone H3 (H3). Control blots targeting the C-terminal end of histone H3 (H3) were included in all WB experiments to ensure equivalent loading between the conditions. Each column represents a different HeLa cell chromatin extract preparation (HeLa #1-4). Each line represents a cropped gel/blot picture (upper/lower marker: 20kDa-15kDa) of several anti-Histone PTMs western blots. The quantitative analysis of the Western blot results is shown alongside in Supporting Figure 2. **C.** Quantification of histone PTMs using specific Nu.Q^®^ immunoassays. Each lane represents a different HeLa cell chromatin extract preparation (HeLa #1-4). The heat map illustrates the relative quantification of each PTM normalized to the H3.1 quantification (%).

**Figure 3.**
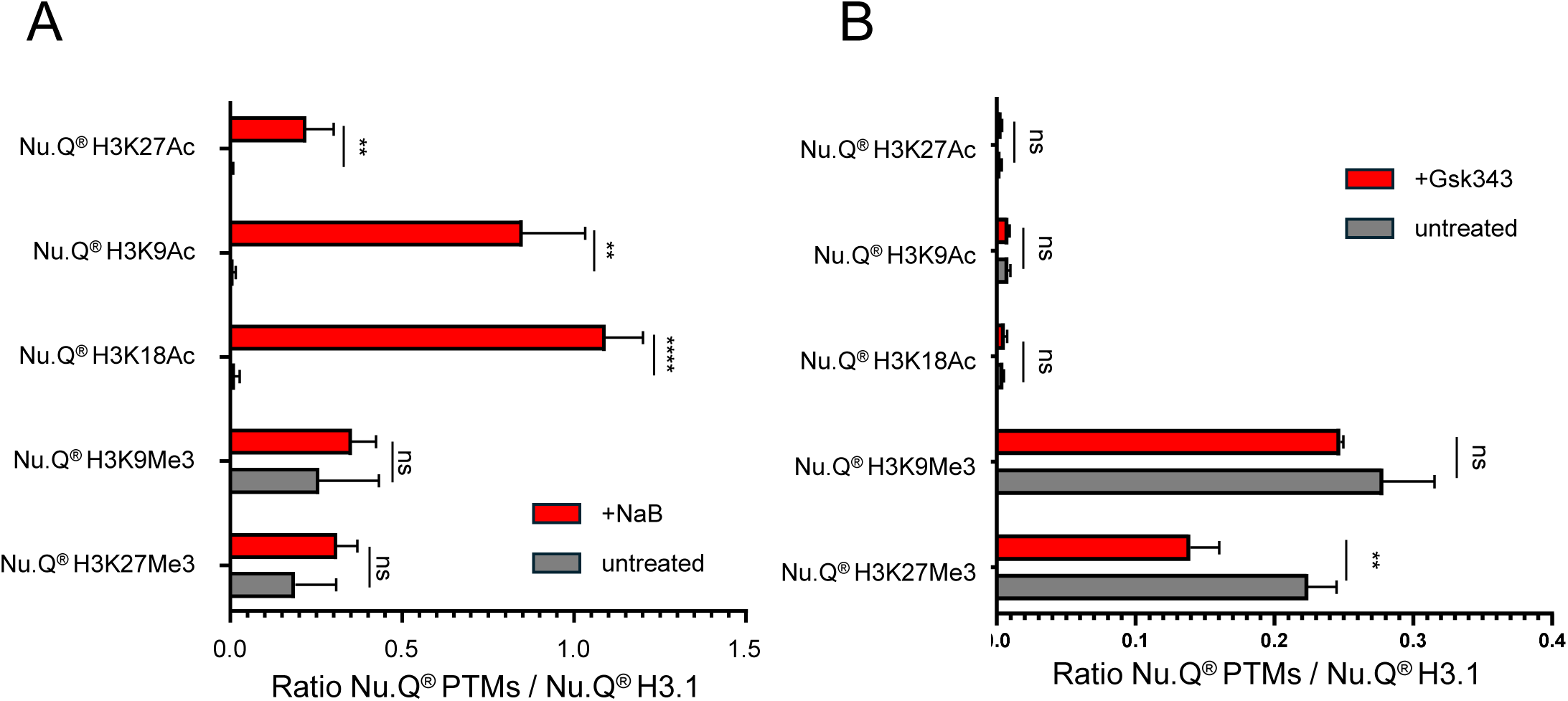
Evaluation of Histone H3 PTMs in HeLa cells treated with Epigenetic modifiers such as Sodium Butyrate (NaB) or EZH2 inhibitor (GSK343). Quantification of Histone H3 Acetylation (H3K27Ac, H3K9Ac, H3K18Ac) and Methylation Levels (H3K9Me3, H3K27Me3) using Nu.Q^®^ Immunoassays (ratio of assay-specific PTMs to the Nu.Q^®^ H3.1 level) in HeLa cells treated with NaB (Red) vs untreated cells (Grey) **(A)** or treated with GSK343, an EZH2 inhibitor (red) vs untreated cells (Grey) **(B)**. Each bar represents the average PTM ratio levels across the three replicates, with error bars indicating standard deviation (SD)(n= 3, Multiple unpaired t-test; \*\**p*□<□0.01; \*\*\*\**p*□<□0.0001; ns: not significant).

### Effective histone PTMs analysis using Nu.Q^®^PTMs assays on chromatin extracted from human fresh-frozen tissues

To validate the applicability of Nu.Q^®^PTMs assays beyond plasma and cellular extracts, we examined nucleosome modification patterns in fresh-frozen human tissues from various organs. First, we confirmed that our optimized custom chromatin extraction protocol produces consistent mononucleosome extracts (∼150bp) across different tissue types, as demonstrated by the comparable DNA profiles obtained using a bioanalyzer (Figure 4A and Supporting Figure 3). Subsequent characterization of histone PTMs/mutations marks revealed a range of modification levels across the tissues. High to moderate levels were observed for H3K27Me3-(range from 11.33% to 95.13%) and H3K36Me3-nucleosomes (range from 13.37% to 173.90%). Moderate to low levels were found for H3R8Cit-(range from 0.03% to 11.77%), H3K27Ac-(range from 0.12% to 9.31%) and H3K9Me3-nucleosome (range from 0.67% to 17.19%). Low levels were detected for H3K4Me2-(range from 0.98% to 6.03%), H3K9Me1-(range from 0.37% to 5.58%), H3K4Me1-(range from 0.38% to 3.19%), H3K18Ac-(range from 0.2% to 4.45%), H3K9Ac-nucleosomes (range from 0.46% to 2.31%) (Figure 4B). In addition, the H3K27M, pH2Ax and H3S10ph-nucleosome markers were not detected in any of the tissues (Data not shown). Interestingly, we observed considerable heterogeneity in the epigenetic patterns across different tissue types and patients, with coefficients of variation ranging from 45% for H3K9Ac-nucleosomes to 226% for H3K27Ac-nucleosomes. Together, these findings suggest that the Nu.Q^®^PTMs assays are effective in characterizing histone PTMs marks across various human tissues.

**Figure 4.**
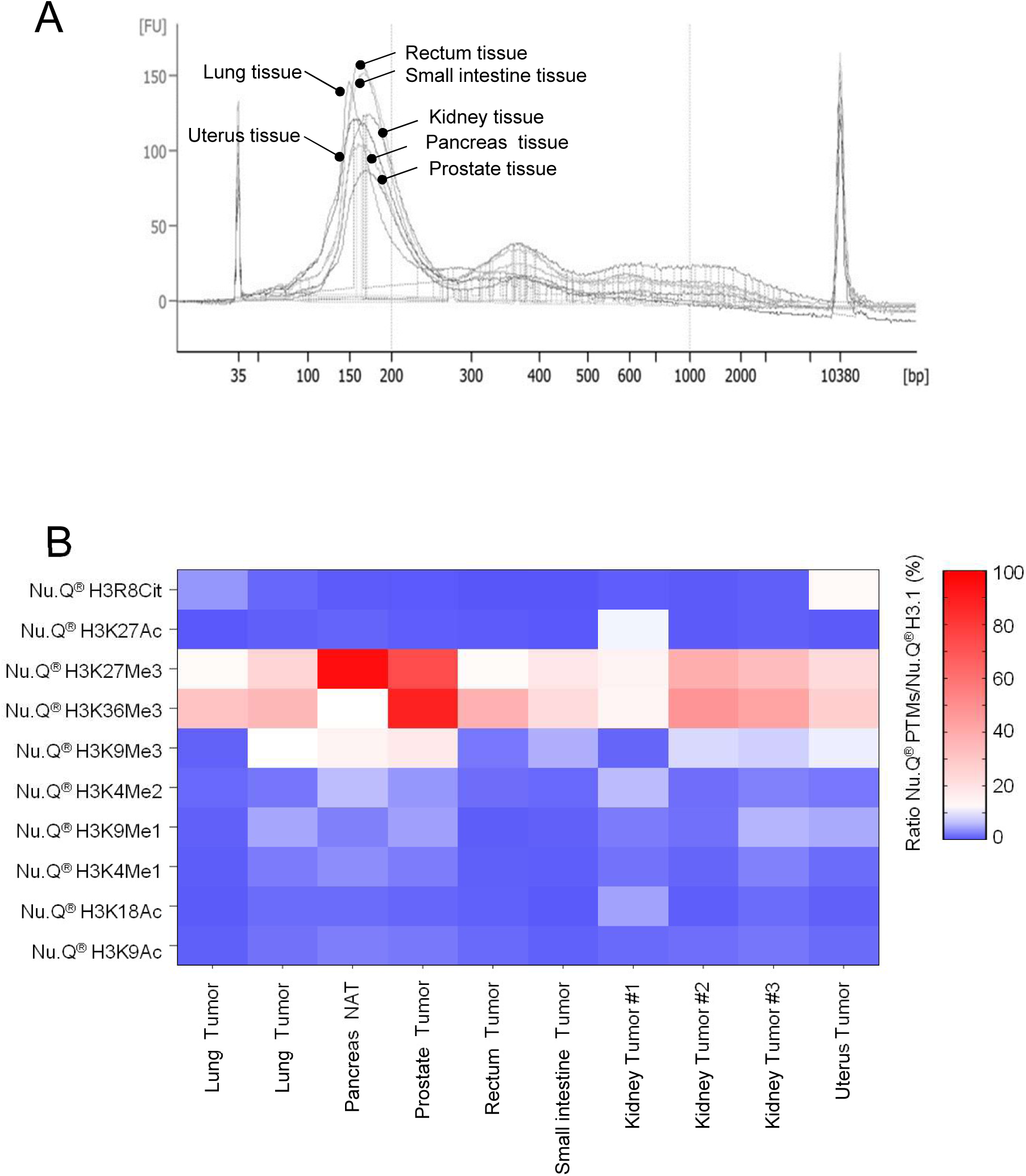
Nucleosome modification patterns in fresh-frozen human tissues from several organ origins. **A.** dsDNA Fragment size distribution of representative chromatin extract from several tissue origine using Agilent 2100 Bioanalyzer. Additional electropherograms are shown in Supporting Figure 3. **B.** The heat map illustrates the relative quantification (%) of each PTM normalized to the H3.1 quantification (lines) in chromatin extracts from different human tissue types (column) including Tumor tissues and Normal adjacent tissues (NAT).

### Histone PTMs insights from subpopulation of white blood cells enabled by cutting-edge Nu.Q^®^PTMs assays

To explore the broader utility of our Nu.Q^®^PTMs assays, we investigated epigenetic patterns in various white blood cell subpopulations. We began by determining the minimal cell count required for efficient chromatin extraction from human peripheral blood mononuclear cells (PBMCs) by testing three different cell quantities (1; 2 and 5 million cells). The DNA profiles obtained from these extractions were analyzed using a bioanalyzer, which consistently confirmed the presence of a mononucleosome peak (∼150bp) in all cases (Figure 5A). Following this confirmation, we measured the amount of H3.1-nucleosomes using the Nu.Q^®^ H3.1 assay and observed a proportional increase in nucleosome quantity with higher cell counts (Figure 5B). Next, we profiled the histone PTMs pattern in PBMCs, monocyte subsets isolated from PBMCs and the non-isolated residual cells (referred to as Monocyte-depleted PBMCs) using the panel of 13 Nu.Q^®^PTMs assays. Subsequent characterization of histone PTMs marks revealed a range of modification levels across the white blood cells subtype. High to moderate levels were observed for H3K9Me3-, H3K27Me3- and H3K36Me3-nucleosomes in all white blood cells subpopulation (n=3; Range observed in PBMCs vs Monocytes vs Monocyte-depleted PBMCs; H3K9Me3- = 0.36% to 22.48% vs 19.59% to 44.81% vs 2.11% to 37.42%; H3K27Me3- = 8.13% to 23.51% vs 16.22% to 28.27% vs 21.06% to 28.61%; H3K36Me3- = 28.01% to 34.01% vs 23.48% to 26.43% vs 17.62% to 39.37%-nucleosomes). Moderate to low levels were found for H3R8Cit-, H3K27Ac-, H3K4Me1-, H3K4Me2- and H3K9Me1-nucleosome (n=3; Range observed in PBMCs vs Monocytes vs Monocyte-depleted PBMCs; H3R8Cit- = 0.64% to 2.39% vs 1.74% to 5.06% vs 0.81 to 3.01%, H3K27Ac- = 0.16% to 0.63% vs 0.27% to 0.98% vs 0.16% to 16.81%; H3K4Me1- = 0.37% to 6.63% vs 3.76% to 9.93% vs 0.56% to 9.41%; H3K4Me2- = 0.35% to 1.16% vs 1.84% to 2.40% vs 1.37% to 11.99%; H3K9Me1- = 0.21% to 12.35% vs 15.36% to 26.38% vs 0.91% to 20.65%-nucleosomes). Low levels were detected for H3K18Ac (range from 0.29% to 1.85%-nucleosomes) and H3K9Ac-nucleosomes (range from 0.00% to 1.88%-nucleosomes). (Figure 5C). In addition, the Nu.Q^®^ H3K27M, Nu.Q^®^ pH2Ax and Nu.Q^®^ H3S10ph markers were not detected in any of the cell extracts (Data not shown). Together, these data suggest that the Nu.Q^®^PTMs assays effectively characterize histone PTMs marks across different subpopulations of white blood cells.

**Figure 5.**
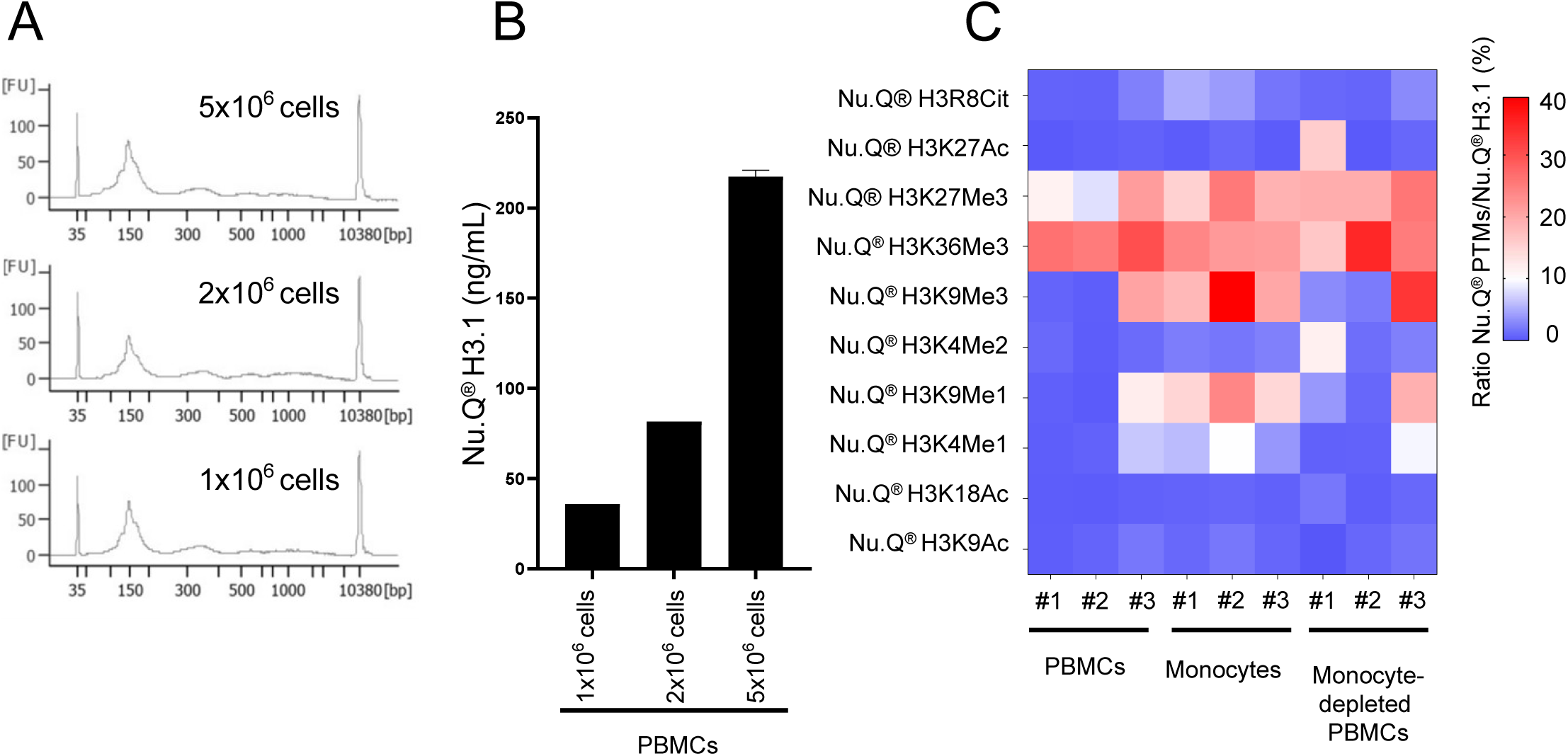
Histone PTMs profiling in white blood cell subpopulations using Nu.Q^®^ assays. **A.** dsDNA Fragment size distribution of dsDNA chromatin extract from multiple number (i.e. 1×10^6^, 2×10^6^, 5×10^6^) of extracted PBMCs cells using Agilent 2100 Bioanalyzer. **B.** Histogram showing the proportional increase in H3.1-nucleosome levels as measured by the Nu.Q^®^ H3.1 assay across increasing PBMC counts (from 1×10^6^ to 5×10^6^ cells). **C.** Heat map illustrating the relative quantification (%), of each histone PTMs normalized to the H3.1 quantification, from PBMCs, isolated monocyte subsets, and the remaining non-isolated white blood cells (referred to as Monocyte-depleted PBMCs).

## Discussion

Epigenetics is a promising field for patient care, offering potential breakthroughs in the diagnosis, identification of novel therapeutic targets and improved treatment monitoring of a wide range of diseases (14, 49, 50). Histone post-translational modifications represent a crucial epigenetic mechanism with strong diagnostic potential (49, 51) as abnormal histone-PTMs patterns are frequently linked to a range of diseases, such as cancer, neurological disorders, and inflammatory conditions (13, 14, 21, 22, 52–56). To study these epigenetic changes, particularly histone-PTMs, several advanced research methods are used such as ChIP-Seq/ChIP-PCR, MS or WB (57–62). In this study, we propose to use sandwich immunoassays to quantify specific histone-PTMs associated with nucleosomes as new powerful tools for epigenetic characterization of circulating nucleosomes, as well as chromatin extraction from cell lines to white blood cells subpopulations and human tissues. Using cell line extracts, we first compared the WB analysis of anti-histone PTMs and MS data to the Nu.Q^®^ immunoassay results, observing a similar epigenetic pattern across all three methods. MS has emerged as a powerful tool for the unbiased and quantitative evaluation of proteins and their PTMs (44, 63, 64). However, this technique involves a complicated process to reduce the sample’s protein complexity such as enrichment of nucleosomes (47, 65) or fractionation using SDS-PAGE gel or ultra-HPLC separation (66), which can potentially lead to the loss of sensitive PTMs (67) (67) or difficulty in detecting certain histone peptides due to their low abundance or structural constraints (68)Additionally, the pretreatment required to handle high lysine content and reduce interfering reagents like high salt or detergents (67, 69, 70) combined with the presence of isobaric peptides (57) can complicate the accurate assignment of PTMs and make relative quantification of PTMs more challenging (44). The ambiguity in the PTM assignation arise when isobaric PTM combinations produce identical mass shifts, mimicking modifications in histone variants. Consequently, this overlap can generate isobaric peptides, making the accurate assignment of PTMs even more challenging (71). As a result, MS often requires validation by another technique, such as WB or immunoassays to ensure accurate identification and quantification of PTMs. WB is widely used to study histone PTMs, but its lack of high-throughput capability and limited linear range make analyzing multiple PTMs costly, labor-intensive, and less accurate (70–72).In addition, the differences in antibody sensitivity hinder the accurate comparison of various PTMs. In contrast, immunoassays are commonly used to evaluate the levels of circulating biomarkers for diagnosis, prognosis or treatment follow-up(75–77). Previous studies have specifically focused on the clinical use of Nu.Q^®^ immunoassays in specific diseases, including cancer (24, 47, 65, 78, 79) and sepsis or NETs-related diseases(76, 80, 81). In this study, we present, for the first time, the application of these assays to characterize epigenetic marks across a variety of sample types, including cell lines, white blood cell subpopulations, and fresh/frozen tissues. By detailing the comprehensive analytical performance of the 13 distinct Nu.Q^®^ assays, we demonstrate their potential for epigenetic research. Indeed, following rigorous development and analytical validation, the immunoassay offers several advantages, including high sensitivity, achieved through amplification via a labeling system, high specificity, shorter turnaround times, simpler procedures, and automation that enables to high-throughput capability. These characteristics make immunoassays highly specific and sensitive, rendering them the methods of choice for measuring complex molecules (82, 83). In this context, Nu.Q^®^ assay offers a promising tool to detect and quantified Histone PTMs in biological samples. Although, the current developed Nu.Q^®^ Immunoassays target 13 different epigenetic marks, mainly localized to the N-terminal region of the histone H3, it does not yet encompass all nucleosome-associated epigenetic modifications, such as crotonylation, lactylation, or ubiquitination, nor does it cover histone post-translational modifications (PTMs) on other histone proteins, like histone H4 (e.g., H4K16Me3). Therefore, alternative technologies may be better suited to identify new histone PTMs that have yet to be investigated. Nevertheless, these assays are highly sensitive in detecting specific histone post-translational modifications associated with circulating nucleosomes. Moreover, these assays are designed to be simple, low-cost and easily accessible, requiring only a minimal sample volume of 50µl. They offer the advantages of automation and rapid processing, delivering reliable results in less than one hour, making them suitable for routine testing. By applying validated immunoassays across a variety of biological samples, ranging from human fresh frozen tissues to specific white blood cell subtypes, we demonstrated the versatility of these assays in epigenetic profiling. For that purpose, a custom in-house extraction protocol was applied across all cells and fresh frozen tissues, yielding satisfactory chromatin extracts. However, this protocol was ineffective on formalin-fixed paraffin-embedded (FFPE) tissues. In addition, an optimization of the extraction protocol may be necessary to analyze more labile marks, like phosphorylation, more effectively. Moreover, given that histone PTMs can have opposite effects depending on the tumor type (50, 84, 85) and that sample sizes for each disease or cell subpopulation in this study are limited, more comprehensive clinical investigations and tissue-specific analyses are essential to identify potential biomarkers and therapeutic targets. Nevertheless, these tools have the potential to contribute to disease research and clinical development, offering a useful resource in epigenetics.

In summary, our work underscores the potential of Nu.Q^®^ immunoassays for precise quantification of PTMs-nucleosomes, showing that analytically validated assays mark a significant step forward in clinical epigenetics research.

## Material and method

### Nucleosomes quantification

H3.1-Nucleosomes level (Nu.Q^®^ H3.1) and specific PTMs-nucleosomes were quantified by Nu.Q^®^ immunoassays (Nu.Q^®^ -H3K27Me3, -H3K36Me3, -H3K4Me2, -H3K9Me3, -H3K9Me1, -H3K4Me1, -H3K18Ac, -H3K9Ac, -H3K27Ac, -H3K27M, -H3R8Cit; -ph2Ax, -H3S10ph) according to the manufacturer’s instructions (Belgian Volition SRL, Isnes, Belgium), after dilution of chromatin extract in a plasma-diluent (Seracon II Delipidated, Seracare; #1800-0017). All immunoassays were developed on the IDS-i10 automated chemiluminescence immunoanalyzer system (Immunodiagnostic Systems Ltd; Boldon, UK) utilizing magnetic beads technology. Briefly, the process began with mixing 50 μL of samples with an acridinium ester labeled anti-nucleosome detection antibody, followed by an incubation period of 30 minutes. Next, magnetic beads coated with either anti-histone H3.1 or histone PTMs antibody were introduced in the reaction and incubated for an additional 15 minutes. Unbound components were then eliminated through a washing step. Finally, trigger solutions were added, and the resulting luminescence was measured with a luminometer. Calibration was achieved using an assay-specific standard curve, and sample quantification was conducted via four-parameter logistic curve fitting.

### Analytical performances

*Sensitivity limits* were established following the recommendations in CLSI guideline EP17-A2 with some modifications. Briefly, to determine the Limit of Blank (LOB), >10 diluent measurements over 5 days were averaged, with the LOB set as the average plus 1.645 standard deviations. The Limit of Detection (LOD) was determined by testing non-blank samples of low concentrations over five days, for a minimum of 30 measurements overall. The pooled standard deviation for all measurements was calculated, and the LOD was determined as LOD = LOB + 1.645 times the pooled SD of non-blank samples. Finally, the lower limit of quantification (LOQ) was defined using a precision profile, with a maximum allowable imprecision of 20% CV for precise quantification.

*Precision* was evaluated via measurement of 4-6 samples displaying different concentration levels, after spiking on human K2EDTA plasma samples with native nucleosomes isolated from human cells or recombinant nucleosomes. The spiked samples were quantified across a minimum of 5 days, with two runs per day and 3 measurements per run. CV% were determined based on CLSI-EP05A guidelines, using an “Analysis of variance” with the Analyse-it software, to assess within-lot and within-laboratory precision. The table presented in the main Figure 1B shows the min to max %CV obtained for each Nu.Q^®^PTMs Assay.

To assess linearity, human K2EDTA plasma samples of varying concentrations were blended in equal proportions (at 10% intervals) and linear curve fitting with variance-based weighting were applied. The linear ranges were defined through comparison of measured concentrations of the different sample proportions with linear regression expected concentrations. The linearity range specifies the interval where the deviation between actual measurements and predictions from the linear model is under 10%. Figure 1D shows a visual representation of linear ranges found on each immunoassay for 1 representative sample. The R², also called Pearson’s linear coefficient of determination, measures the strength of the linear relationship between the response variable (Y) and the predictor variable (X) on the linearity range determined. This value stands between 0 and 1 and therefore makes it possible to evaluate the relevance of the linear model for the dilutions of our samples.

*Specificity:* The cross-reactivity was assessed by quantifying single recombinant nucleosome (rNucl.; (Belgian Volition SRL, Isnes, Belgium)) exhibiting a single, precisely localized epigenetic mark such as either a variant modification (H3.1 or H3.3 histone) or PTMs or Mutation (H3K27M) using the Nu.Q^®^ assay panel. All Nu.Q^®^ tests were conducted simultaneously using the same rNucl., diluted at 1µg/mL in a plasma diluent (Seracon II Delipidated, Seracare; #1800-0017) preparation.

### Samples and cells collection

Frozen human PBMCs, diluted in 60% RPMI-1640, 35% FBS and 5% DMSO, and fresh-frozen human tissues were purchased from Biomedica CRO, a biobank that presents a full compliance with the international and local ethical guidelines (approval by the IRB/IEC). HeLa S3 cells (ATCC, #CCL-2.2) were cultured in S-MEM medium (Gibco, #11380037) supplemented with 10% Fetal Bovine Serum (Gibco, #A4766801), 1x NEAA (Non-Essential Amino Acids solution) (Gibco, #11140-050), Penicillin/Streptomycin (Gibco, #15070063) and 1x L-glutamine (Gibco, #25030081), with or without NaB treatment (5mM as final concentration, for 20h; Merck Life Sciences BV; #19-137) or with / without GSK343 treatment (5µM as final concentration, for 48h; Abcam, #Ab218610). HeLa S3 cells were maintained in a CO_2_ incubator (37°C, 5% CO_2_).

### Monocytes isolation

After evaluating the number of frozen PBMCs cells using the NanoEnTek, EVE TM Plus system, the cells were centrifuged at 200g for 10 minutes, and the cell pellet was resuspended on 1mL of isolation buffer (phosphate-buffered saline (PBS), pH 7.2, 0.5% bovine serum albumin (BSA) (VWR; #4215015), 2mM EDTA (Merck Life Sciences BV; #E7889)) and recentrifuged to pellet PBMCs. Monocytes were then isolated following Pan-Monocyte Isolation Kit instructions (Miltenyi; 130-096-537).

### Chromatin extraction

*From cells (i.e. PBMCs, monocytes and/or HeLa cells)*: After cell counting (NanoEnTek; EVE TM Plus), cells were centrifuged for 5 minutes at 150 g and washed with 1mL of cold PBS. For each 2.5 ×10^7^ cells, cells pellets were resuspended on 300µl of lysis buffer (10mM HEPES pH 7.4, 10mM KCL, 1mM Sodium orthovanadate, 0.05 % Igepal (Merck Life Sciences BV; #i8896), EDTA-free cOmplete^TM^ (Roche; #4693159001)) and incubated for 20 minutes on ice before being centrifuged again at 12,000 g for 10 minutes. This was followed by an additional wash cycle involving a lysis buffer, incubation, and centrifugation steps. Cells pellet was then resuspended on 600µL of low salt buffer (10mM Tris-HCl pH 7.4, 0.2mM MgCl2, 1mM Sodium orthovanadate, 1% Triton (Merck Life Sciences BV; #T8787) + EDTA-free cOmplete^TM^ (Roche; #4693159001)) and incubated on ice for 15 minutes before being centrifuged again at 12,000 g for 10 minutes. Then, the cells pellet was resuspended on 100µL MNAse digestion buffer (5mM CaCl2, 50mM Tris-HCl pH 8, 1mM sodium orthovanadate + EDTA-free cOmplete^TM^ (Roche; #4693159001)). Finally, 0.155U of MNAse (Merck Life Sciences BV; #N5386) was added to each sample, and incubate at 37°C for 15 min. After that step, 50µL of EDTA 0.5M (Merck Life Sciences BV; #E7889-100ml) were added to the mixture and incubate 5 minutes on ice to stop MNAse reaction. Nucleosomes were then recovered on supernatant after centrifugation at 9,000g for 5 minutes. For NaB treated HeLa S3 treated, 5mM of NaB (Merck Life Sciences BV; #19-137) was added to all extraction buffers (i.e. Lysis buffer, Low salt buffer and MNAse digestion buffer).

*From Fresh Frozen tissues:* 100mg of frozen tissues were fixed with 40µL of 1% PBS – formaldehyde (FA-PBS) (Merck Life Sciences BV; #F8775) and then finely chopped with little scissors. Then, 900μL of 1% FA-PBS was added to the tissues and the tissues were fixed for 10 minutes at RT on a rotating wheel. Subsequently, 0.125M of Glycine (Cytiva; #GE17-1323-01) was added and incubated for an additional 10 minutes on a rotating wheel. Following this, a wash (500µL of PBS + 1% of protease inhibitor cocktail (Merck Life Sciences BV; #P8340)) and centrifugation cycle (12,000g for 2 min) was performed three times consecutively. After that, the fixed tissues were incubated with either 1mL of 1% collagenase/Dispase-PBS (Roche; 10269638001) containing 0.1% of CaCl2 1M (Merck Life Sciences BV; #C1016-500g) for 30 minutes at 37°C (i.e. Rectum, pancreas and small intestine tissues) or 500µL of trypsin-EDTA (Thermo Fisher Scientific; 25300-054) for 10 min at 37°C (i.e. Kidney, lung, uterus and prostate tissues). Tubes were then centrifuged at 12,000g for 5 minutes and supernatant was discarded. A last wash with PBS-1% protease inhibitor was performed. Tissues pellet was resuspended on 360µL of lysis buffer (10mM HEPES pH 7.4, 10mM KCL, 1mM Sodium orthovanadate, 0.05 % Igepal, EDTA-free cOmplete^TM^) and mechanical disruption was performed using a homogenizer with a pestle adapter (BELART; #651000000) followed by an incubation on ice for 20 min. After a centrifugation at 12,000g for 10 minutes and a wash step with lysis buffer, tissue pellets were then resuspended on 720µL of low salt buffer (10mM Tris-HCl pH 7.4; 0.2mM MgCl2, 1mM Sodium orthovanadate, 1% Triton, EDTA-free cOmplete^TM^). After an incubation of 15 minutes on ice and a centrifugation at 12,000g for 10 minutes, tissues pellets were resuspended on 120µL MNAse digestion buffer, 0.2U of MNAse (Merck Life Sciences BV; #N5386) were added to each sample, and incubate at 37°C for 30 min. After that step, 12µL of EDTA 0.5M (Merck Life Sciences BV; #E7889) were added to the mixture and incubate 5 minutes on ice to stop MNAse reaction. Nucleosomes were then recovered on supernatant after centrifugation at 9,000g for 5 minutes.

### Characterization of Histone post-translational modifications by Liquid chromatography with tandem mass spectrometry (LC–MS/MS)

HeLa cells nucleosomes were extracted following previously described protocol, and were then processed by EpiQMAx GmbH (Planneg, Germany) to analyze the histone post-translational modifications pattern by LC–MS/MS. Nucleosomes were propionylated and trypsinized. Heavy amino acid-labeled Histone H3 peptides were added to digested samples for normalization and quantifications purposes. The resulting mixture was separated by liquid chromatography (Ultimate 3000 RSLC nano System (Thermo-Fisher Scientific) containing a 15-cm analytical column (75 μm ID with ReproSil-Pur C18-AQ 2.4 μm from Dr. Maisch) directly coupled to an electrospray ionization source and Q Exactive HF mass spectrometer (Thermo-Fisher Scientific). A data-dependent acquisition mode was used to automatically switch between full scan MS and MS/MS acquisition. For the analysis, the raw peptides of canonical histone peptides and their respective PTMs intensities were searched with Skyline software and normalized using the intensities of corresponding spiked-in heavy peptides. After normalizing the light peptides using heavy peptides, we presented the mass spectrometry (MS) results as log_₂_-transformed normalized intensities.

### Western Blotting

*Experimental process :* a total of 200ng of four different chromatin extracts from HeLa cells (#1-#4) were loaded onto a Western blot gel, along with a molecular weight ladder and a specific histone PTMs recombinant nucleosome (r.Nucl) used as a positive control. Chromatin extract were mixed with Laemmli buffer (Biorad; #1610747) containing 10% of 2-mercaptoethanol (Merck Life Sciences BV; #M6250) and denaturated for 5 minutes at 95°C. Denatured nucleosomes were separated by electrophoresis on 4-20% TGX precast gels (Biorad; #4561094) under reducing conditions (Tris/Glycine/SDS; Biorad; #1610772) and then transferred to PVDF (polyvinylidene fluoride) membranes using a Trans-Blot Turbo semi-dry transfer system (Biorad; #1704272). The PVDF membranes was then incubated 2 hours with antibodies against H3K27Me3 (Cell Signaling Technology; #9733BF – lot: 25), H3K36Me3 (Active Motif; #80218 – lot: 2280106), H3K9Me3 (Abcam; #232324 – lot: GR3452452-1), H3K4Me2 (Volition; #3349 lot: rr260604a-6240), H3R8Cit (Abcam; #232939 – lot: GR3402039-3), H3K27Ac (Volition; #3118– lot: rr200819b-6298), H3K18Ac (Synabs; #FYN213R CL3 lot: 10487), H3K9Ac (Active Motif; #91103 – lot: 22420006), H3K9Me1 (Cell Signaling Technology; #14186BF – lot: 5), H3K4Me1 (Cell Signaling Technology; #5326BF – lot: 4), diluted on Tris Buffer Saline, 1% of Casein (Biorad; #1706435) + 0.1% Tween^®^20 (Merck Life Sciences BV; #T2700) at 2µg/ml and incubated for 2H at RT or overnight at 4°C. After primary antibody incubation, membranes were washed three times with 1x TBST ((Biorad; #1706435) + 0.1% Tween^®^20 (Merck Life Sciences BV; #T2700)) and incubated with a detection reagent (VeriBlot antibody for IP Detection Reagent, Abcam; #ab131366) or anti-rat HRP antibody (for H3K18AC detection; Jackson immunoresearch, 112-035-003 – lot: 146459) for 1 hour at RT. Membranes were again washed three times with TBST and incubated with Super Signal West Dura Extended Duration Substrate for 5 minutes (Life Technologies Europe, #34076). Chemiluminescent signals were acquired with the Fusion-FX6 instrument and software (Vilber). An anti-H3 C-terminal Western blot was then performed on all membranes after a stripping step to remove detection antibodies. For that purpose, membranes were incubated twice for 10 minutes with a mild stripping buffer (Glycine 0.2M, SDS 0.1%, Tween^®^20 1%, pH 2.2), followed by two washes on 1x PBS and 2 washes on 1x TBST. Stripped membranes were then reincubated with anti-H3 C-terminal antibody (Active Motif; #91297 – lot: 316202) diluted at 1.5µg/ml on TBS, 1% of Casein (Biorad; #1706435) + 0.1% Tween^®^20 (Merck Life Sciences BV; #T2700). The following steps were the same as previously described for all detection antibodies. To validate the specificity of the detected signals, recombinant nucleosomes specific to each PTMs were loaded on a separate lane (data not shown).

Western-blots were quantified using Bio-1D quantification program (Vilber Lourmat) and normalized based on associate H3C-terminal band signal. Quantification of the specific histone-PTMs in the sample was defined by comparing the band intensity of sample versus the band intensity of the known quantity of the positive control. Then, Western blots targeting post-translational modifications were normalized to the total level of histone H3 and expressed as a ratio.

### Nucleosomal DNA extraction and DNA size profile

Chromatin extracts were diluted on 1x PBS and extracted with QIAamp^®^ DSP Circulating NA kit (Qiagen; #61504) following manufacturer’s instructions. The subsequent DNA size distribution was analyzed by using Agilent High Sensitivity DNA Kit for fragment sizes of 50 - 7000bp (Agilent Technologies; #5067-4626) with the Agilent 2100 Bioanalyzer (Agilent Technologies), following the manufacturer’s instructions.

### Statistical analysis

*Heat map analysis* were performed using GraphPad InStat software (GraphPad Software, USA). Mass spectrometry data are represented as Log_2_ of normalized intensity. Nu.Q^®^ PTMs are represented as a ratio (%) of Nu.Q^®^ PTMs/Nu.Q^®^ H3.1. Specificity data are represented as the % of cross-reactivity. The cross-reactivity was measured and calculated by dividing the concentration of rNucl. exhibiting unspecific PTMs by the concentration obtained for rNucl. exhibiting specific PTMs (%).

Multiple unpaired t-tests, using GraphPad InStat software (GraphPad Software, USA), were conducted to evaluate the effect of NaB and GSK343 on the epigenetic profile in HeLa cells \*\**p*□<□0.01; \*\*\*\**p*□<□0.0001).

## List of abbreviations

Ac: Acetyl
bp: Base pair
BSA: Bovine serum albumin
ChIP: Chromatin immunoprecipitation
ChLIA: Chemiluminescent immunoassays
Cit: Citrullination
CV: Coefficients of variation
dsDNA: double-stranded DNA
FA-PBS: PBS – Formaldehyde
FFPE: Formalin-fixed paraffin-embedded
HDAC: Histone Deacetylase
IHC: Immunohistochemistry
K: Lysine
LC-MS/MS: Liquid chromatography with tandem mass spectrometry
LOB: The Limit of Blank
LOD: The Limit of Detection
LOQ: The Limit of Quantification
Me1/Me2/Me3: Methyl ; 2 Methyl ; 3 Methyl
MNAse: Micrococcal nuclease
MS (/ MS): (Tandem) Mass spectrometry
NaB: Sodium butyrate
NAT: Normal Adjacent Tissue
ns: Non-significant
PBMCs: Peripheral blood mononuclear cells
PBS: Phosphate-buffered saline
PCR: Polymerase chain reaction
PTMs: Histone post-translational modifications
rNucl.: Recombinant nucleosomes
RT: Room Temperature
SD: Standard Deviation
SDS-PAGE: Sodium dodecyl-sulfate polyacrylamide gel electrophoresis
Seq: Sequencing
TBST: Tris Buffer Saline + 0.1% Tween® 20
ultra-HPLC: Ultra-High-Performance Liquid Chromatography
WB: Western blotting
WBC: White blood cells

## Acknowledgments

We would like to thank Virginie Laloux and Fanny Lambert for their work in cultivating and amplifying HeLa cells. We thank Moritz Völker-Albert and the EpiQMAx’s team for MS data acquisition and their assistance in drafting the MS material section. We thank the Administrative and Logistics team of Belgian Volition for their support. We thank Louise Batchelor for her careful proofreading of the manuscript. We thank NIAID Visual & Medical Arts from BIOART source for providing the illustration of the antibody (07/10/2024). Antibody. NIAID NIH BIOART Source. bioart.niaid.nih.gov/bioart/17. OpenAI *ChatGPT* GPT-4 was used to support minor edits throughout the manuscript.

## CRediT Author statement

Conceptualization : P.VdA; Analysis and interpretation of data: D.P., C.H., G.R., M.W. and P.VdA.; resources and data acquisition: C.H., M.L., O.T., R.V., S.V. and M.C.; writing original draft: P.VdA, M.W., C.H. and D.P., writing-review and editing D.P. ; C.H., G.R., R.V., M.W., O.T. M.L., S.V., M.C., M.H. and P.VdA. Supervision: M.H. All authors have read and approved the manuscript.

## Additional information: declaration of Interests

All authors are employed by Belgian Volition SRL. Belgian Volition SRL has patents covering Nu.Q^®^ technology and are developers of Nu.Q^®^ assays. Belgian Volition SRL is a subsidiary of VolitionRx Limited. MC and MH are shareholders of VolitionRX Limited.

## Data availability

The datasets used and/or analyzed during the current study are available from the corresponding author on reasonable request.

## Supporting Information

This article contains supporting information.

