## Supporting Figure 1 for "High-Throughput Epigenetic Profiling Immunoassays for Accelerated Disease Research and Clinical Development"

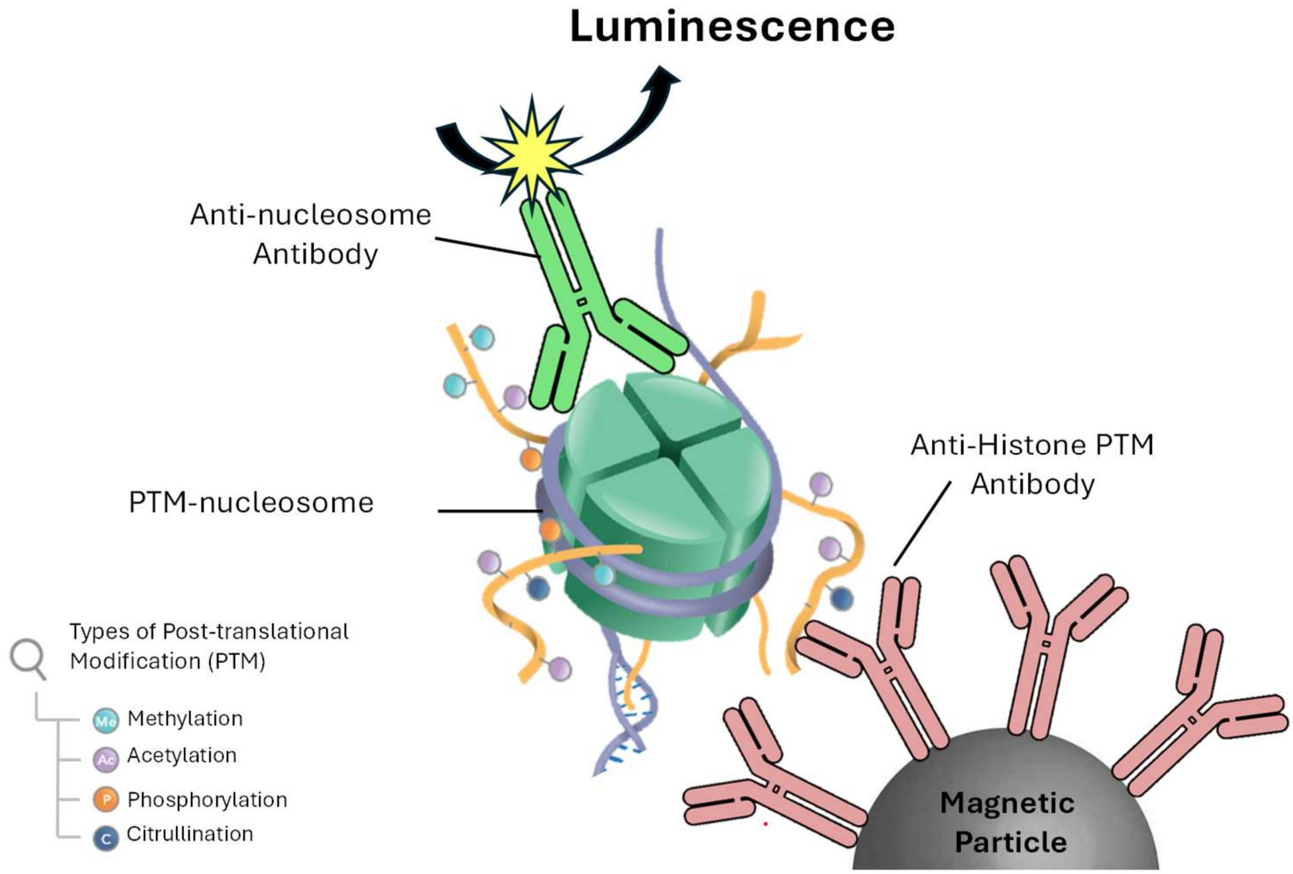


**Supporting Figure 1 :** Visual representation of Nu.Q®immunoassay principles. Nu.Q®immunoassays are sandwich assays using an anti-histone PTMs antibody in capture on the solid phase, consisting of magnetic particles, combined with a conformational anti-nucleosome antibody coupled with acridinium ester molecules in detection.
