## Supporting Figure 2 for "High-Throughput Epigenetic Profiling Immunoassays for Accelerated Disease Research and Clinical Development"

| Membrane | WB anti- | HeLa chromatin extracts | | | |
|---|---|---|---|---|---|
| | | #1 | #2 | #3 | #4 |
| A | *H3K4Me1* | *62,981* | *79,738* | *61,389* | *52,385* |
| | *H3* | *597,202* | *741,801* | *782,129* | *708,925* |
| | **Ratio H3K4Me1/H3** | **0,105** | **0,107** | **0,078** | **0,074** |
| B | *H3K4Me2* | *19,924* | *14,123* | *13,255* | *16,594* |
| | *H3* | *657,750* | *870,216* | *854,774* | *702,462* |
| | **Ratio H3K4Me2/H3** | **0,030** | **0,016** | **0,016** | **0,024** |
| C | *H3K9Me1* | *47,809* | *54,464* | *38,733* | *38,009* |
| | *H3* | *259,726* | *270,049* | *211,638* | *216,307* |
| | **Ratio H3K9Me1/H3** | **0,184** | **0,202** | **0,183** | **0,176** |
| D | *H3K9Me3* | *123,892* | *83,677* | *100,326* | *75,958* |
| | *H3* | *183,574* | *169,439* | *170,222* | *115,246* |
| | **Ratio H3K9Me3/H3** | **0,675** | **0,494** | **0,589** | **0,659** |
| E | *H3K9Ac* | *6,286* | *2,728* | *5,224* | *15,649* |
| | *H3* | *1103,000* | *1352,000* | *1186,000* | *999,162* |
| | **Ratio H3K9Ac/H3** | **0,006** | **0,002** | **0,004** | **0,016** |
| F | *H3K18Ac* | *5,870* | *3,175* | *4,434* | *3,079* |
| | *H3* | *1301,000* | *1236,000* | *1065,000* | *766,473* |
| | **Ratio H3K18Ac/H3** | **0,005** | **0,003** | **0,004** | **0,004** |
| G | *H3K27Me3* | *258,564* | *224,819* | *172,823* | *222,209* |
| | *H3* | *296,004* | *178,858* | *250,660* | *236,019* |
| | **Ratio H3K27Me3/H3** | **0,874** | **1,257** | **0,689** | **0,941** |
| H | *H3K27Ac* | *10,556* | *5,387* | *5,421* | *3,031* |
| | *H3* | *592,444* | *477,060* | *448,709* | *357,677* |
| | **Ratio H3K27Ac/H3** | **0,018** | **0,011** | **0,012** | **0,008** |
| I | *H3K36Me3* | *99,506* | *85,872* | *73,156* | *65,607* |
| | *H3* | *363,931* | *293,644* | *192,121* | *175,895* |
| | **Ratio H3K36Me3/H3** | **0,273** | **0,292** | **0,381** | **0,373** |
| J | *H3R8Cit* | *4,025* | *3,700* | *4,018* | *8,184* |
| | *H3* | *299,042* | *275,450* | *178,772* | *216,814* |
| | **Ratio H3R8Cit/H3** | **0,013** | **0,013** | **0,022** | **0,038** |

**Supporting Figure 2:** The table presents the raw data (expressed in ng) obtained from the quantitative analysis of the Western blot results shown in Figure 2B.
