## Supporting Figure 3 for "High-Throughput Epigenetic Profiling Immunoassays for Accelerated Disease Research and Clinical Development"

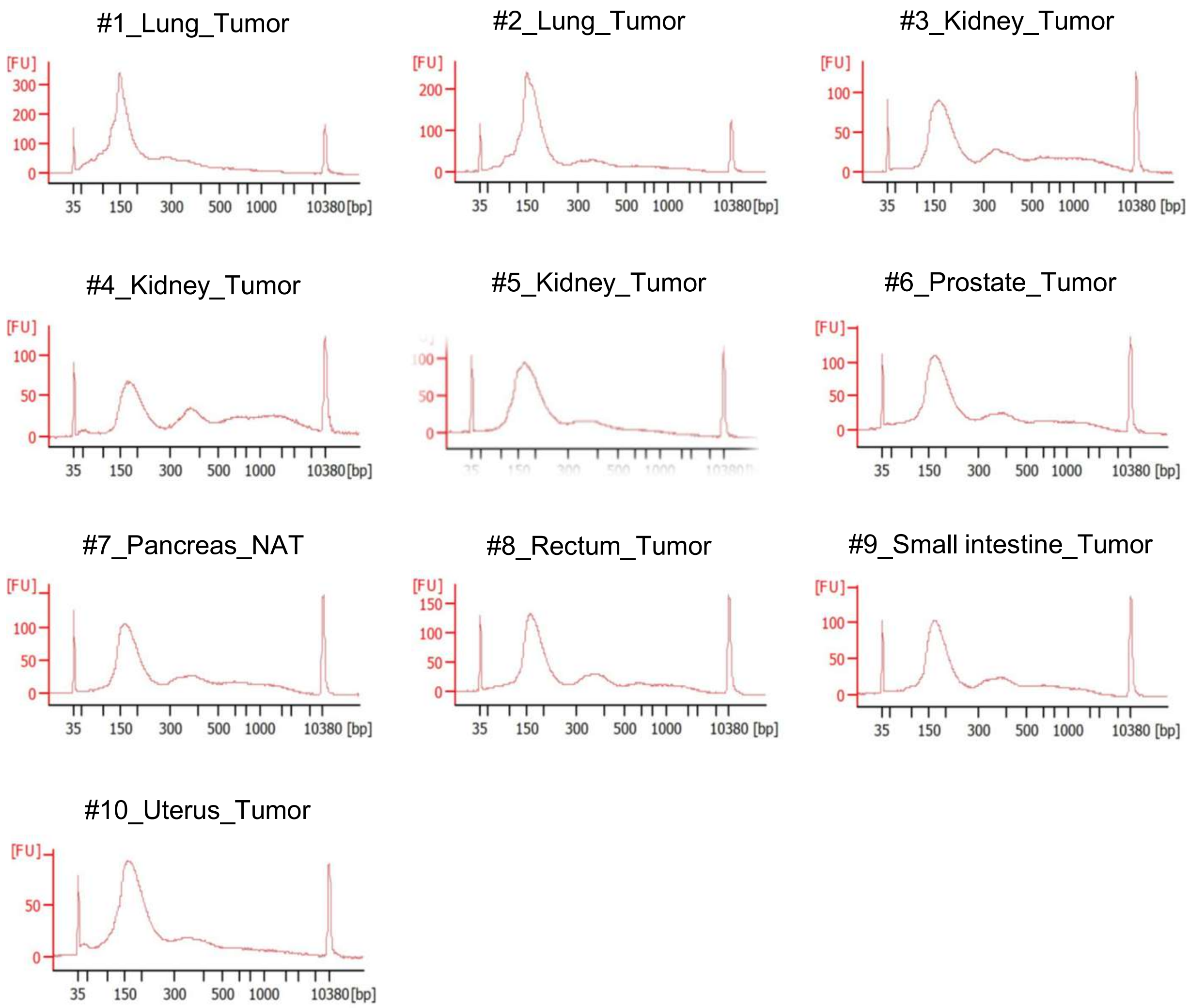


**Supporting Figure 3 :** DNA Fragment size distribution of chromatin extract from several tissue origins shown in Figure 4A using Agilent 2100 Bioanalyzer.
